# Homozygosity at its Limit: Inbreeding Depression in Wild *Arabidopsis arenosa* Populations

**DOI:** 10.1101/2021.01.24.427284

**Authors:** A. Cristina Barragan, Maximilian Collenberg, Rebecca Schwab, Merijn Kerstens, Ilja Bezrukov, Felix Bemm, Doubravka Požárová, Filip Kolář, Detlef Weigel

## Abstract

New combinations of genetic material brought together through hybridization can lead to unfit offspring as a result of outbreeding or inbreeding depression. In selfing plants such as *Arabidopsis thaliana*, outbreeding depression is typically the result of pairwise deleterious epistatic interactions between two alleles that can geographically co-occur. What remains elusive is how often alleles resulting in genetic incompatibilities co-occur in natural populations of outcrossing plant species. To address this question, we screened over two thousand five hundred wild *Arabidopsis arenosa* hybrid plants in search for potential genetic mismatches. We show that although abnormal deleterious phenotypes are common, the transcriptional profiles of these abnormal *A. arenosa* plants differ substantially from those seen in incompatible *A. thaliana* hybrids. The abnormal hybrid phenotypes in *A. arenosa* had different underlying genetic architectures, yet a repeated theme was increased homozygosity, indicating that inbreeding rather than outbreeding depression gives rise to some of the deleterious phenotypes segregating in wild *A. arenosa* populations.

## Introduction

Hybridization resulting in either outbreeding or inbreeding depression can act as a postzygotic reproductive barrier by reducing genetic exchange among individuals^1^. While meiotic hybridization occurs in both selfing and outcrossing species, repeated self-fertilization results in high levels of homozygosity across the genome in selfers. Outcrossers on the other hand, depend on the presence of foreign pollen for fertilization, and are typically more heterozygous^2^. When plants that are typically selfing, outcross, deleterious epistatic interactions between divergent alleles can arise when these are brought together though heterozygosity. This applies to hybrid necrosis cases that follow the Bateson-Dobzhansky-Muller (BDM) model and which have been mainly described in plant species or cultivated variants that are predominantly selfing^3–13^. Alleles underlying incompatibility associated with autoimmunity can co-occur in wild *Arabidopsis thaliana* populations^4,10^. How common such BDM incompatibilities are in local populations of outcrossing species remains unclear.

Selfing plants rapidly purge strongly deleterious alleles, but slightly deleterious mutations tend to accumulate more easily than in outcrossing species^14,15^. Generally, natural selection is expected to eliminate genetic incompatibilities from populations where individuals are frequently interbreeding, unless the advantages incompatible loci confer when present in the same population outweigh the disadvantages caused by their potential incompatibility.

To address these issues, we studied *Arabidopsis arenosa*, an obligate outcrossing relative of *A. thaliana*^16,17^. We investigated eight diploid populations from the Western Carpathian Mountains, which is a center of genetic diversity of the species^18,19^. We collected tissue and corresponding seeds from over 1,700 mother plants, and examined over 2,500 of the naturally hybrid progeny in the lab. We found abnormal hybrid phenotypes to be common and often inherited across generations - more so than in *A. thaliana*^4,9^, but hybrid necrosis appears to be much rarer in *A. arenosa* than its congener *A. thaliana*.

In one family, we identified by linkage mapping a single genomic region that gives rise to its abnormal phenotype when homozygous. In a second family, with a different deleterious phenotype, multiple homozygous regions throughout the genome were shared among abnormal plants. We speculate that various combinations of these regions result in independent deleterious abnormalities. In a third family, although it consistently segregated a chlorotic and dwarf phenotype, we could not identify any obvious QTL or genomic region(s) that was linked to this phenotype, leaving it unclear what the underlying genetics is.

Inbreeding depression is brought on by reducing the advantage some alleles confer when present in a heterozygous state, or by exposing recessive deleterious alleles when in a homozygous state^20^. Our results suggest that it is inbreeding depression, rather than pairwise deleterious allelic interactions, which causes several phenotypic abnormalities segregating in wild *A. arenosa* populations.

## Results

### Structure of Wild *Arabidopsis arenosa* Populations

Plant material and seeds were collected from eight different diploid *A. arenosa* populations in central and western Slovakia (**Fig 1A**, **Table S1**). In each population we sampled, non-destructively, leaf tissue and seeds from the majority of individuals present. The number of sampled plants is therefore proportional to the size of the sampled populations (**Fig S1A**). We collected at least a dozen, but usually a few hundred seeds per plant, at a time when the majority of seeds had already been shed, ensuring the undisturbed persistence of all populations. Seeds from a single mother plant are either siblings (if they share the same pollen donor) or half sibs (if they do not). We will refer to the immediate as well as all later-generation progeny from a single mother plant as ‘families’.

**Fig 1.**
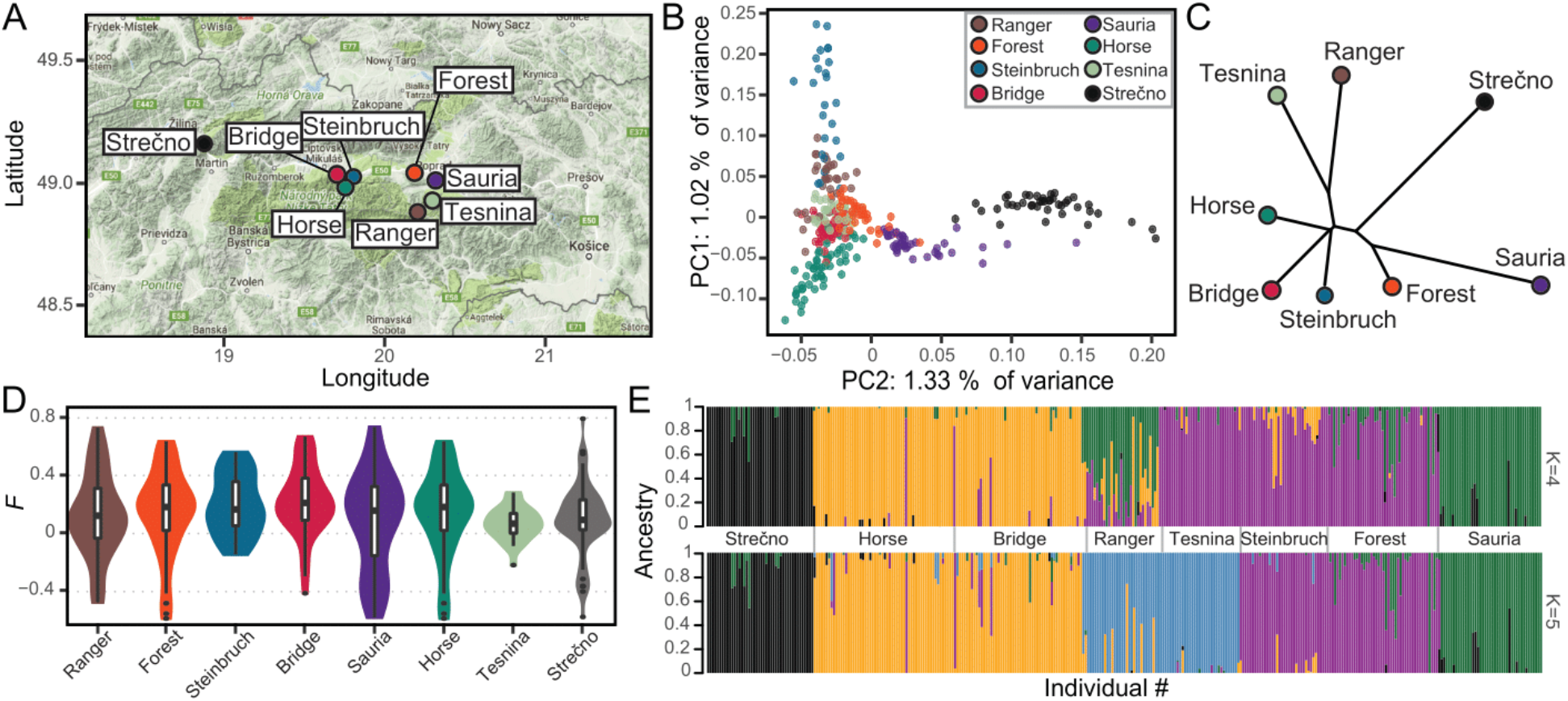
Structure of the sampled *A. arenosa* populations. **A.** Location of the eight sampled *A. arenosa* populations. **B.** Genetic relationship between individual samples estimated by Principal Component Analysis. Variance between populations is explained by multiple small-effect components. **C.** *F*_*ST*_-based Neighbour-Joining (NJ) tree of the eight populations. Isolation by distance is observed. **D.** Distribution of the inbreeding coefficient (*F*) per individual for each population. The average *F* per population ranged from 0.07 to 0.22. **E.** ADMIXTURE plot showing ancestry inferences^24^. Four (K=4) or five (K=5) ancestral populations likely gave rise to the eight populations sampled.

To study the structure of the eight populations, we individually genotyped 345 of the 1,768 sampled mother plants by RAD-seq^21^ (**Table S1**). The sequencing reads were mapped to a newly generated *A. arenosa* chromosome-scale reference genome originating from a plant collected in Strečno, one of the eight populations (**Table S2**). A Principal Component Analysis (PCA) showed that variance between populations is explained by multiple small-effect components (**Fig S1B**), with samples from the same population being genetically more similar to each other (**Fig 1B**, **S1C**). In addition, isolation by distance between populations was observed: the closer two populations are to each other geographically, the more genetically similar they tend to be (**Fig 1C**, **S1D**). The average inbreeding coefficient (*F*) of each population ranged from 0.07 to 0.22 (**Fig 1D**), which is similar to what has been described in populations of the closely related, and also outcrossing, *A. lyrata*^22^. In many cases, a higher apparent *F* value, above 0.5, was explained by a large amount of missing data per individual^23^ (**Fig S1E**). The eight populations analyzed were estimated to have arisen from four or five clearly defined ancestral populations (**Fig 1E**, **S1F**), with populations that are geographically closer tending to share a common ancestor (**Fig 1E**).

### Abnormal and Heritable Phenotypes are Relatively Common in *A. arenosa*

Because *A. arenosa* is an obligate outcrosser, all individuals are by definition hybrids, since they are the result of combining two parental genotypes. To study how prevalent hybrid incompatibilities naturally arise among co-occurring *A. arenosa* individuals, we sowed 5 to 7 seeds from 461 families originating from all eight sampled populations. These plants, which we designate as F_1_ individuals, were screened for abnormal phenotypes. If heritable, these phenotypes could be caused by the presence of deleterious alleles, or by deleterious epistatic interactions between alleles, either at one or multiple loci (**Fig 2A**, **Table S3**).

**Fig 2.**
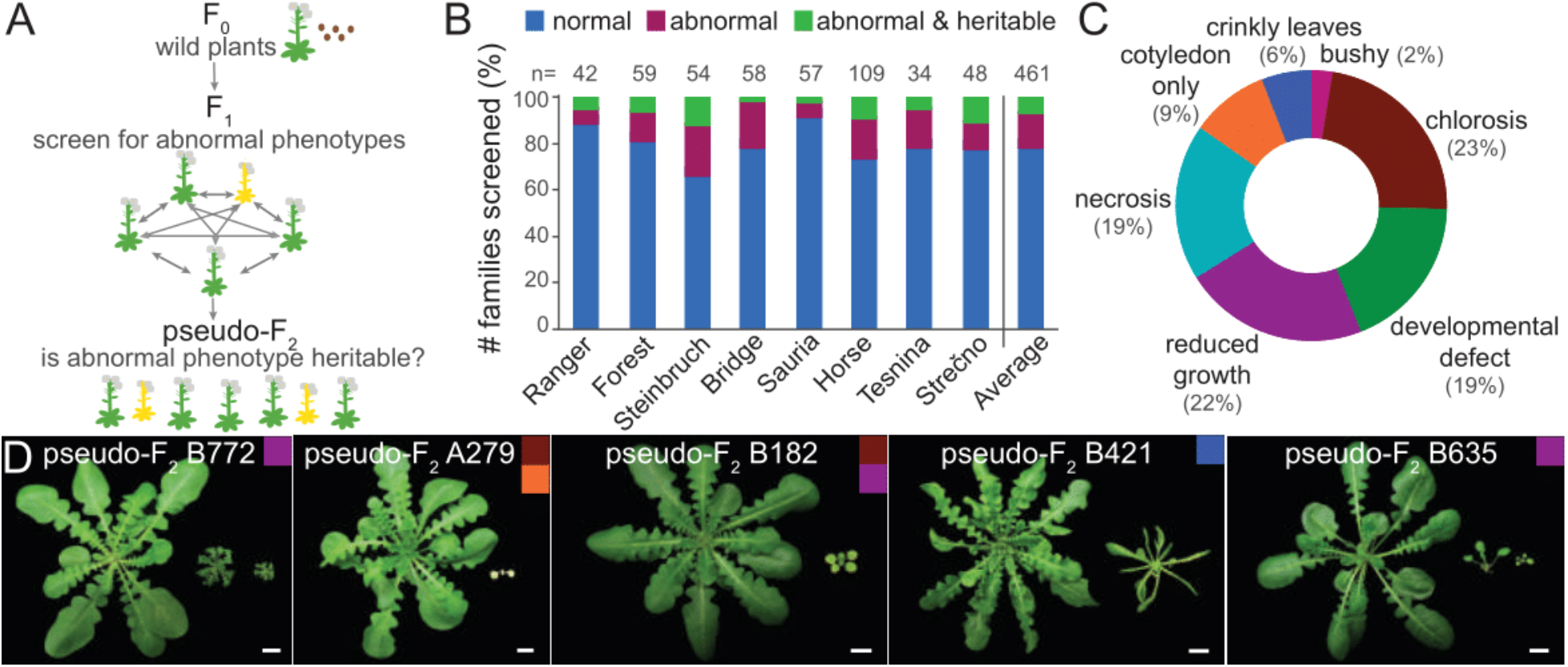
Screening for abnormal *A. arenosa* phenotypes. **A.** Experimental design involving the creation of pseudo-F_2_ populations. F_1_ seeds were collected from wild *A. arenosa* mother plants (F_0_). These F_1_ offspring were screened for abnormal phenotypes in the lab (yellow) and, if present, crossed with their siblings to test whether this phenotype was recapitulated in the following generation (pseudo-F_2_). **B.** Percentages of families showing abnormal phenotypes in the F_1_ generation (abnormal), and the percentage of phenotypes recapitulated in the pseudo-F_2_ generation (abnormal heritable) per geographic population (see Fig 1A). The number of F_1_ families screened per population is indicated at the top (n). **C.** Pie chart showing the most common abnormal phenotypes in the F_1_ plants. Some plants fell under more than one category. **D.** Examples of normal (right) and abnormal and heritable (left) phenotypes from five independent families. Phenotype categories for abnormal plants in each family are indicated, colors as in C. Plants were seven weeks old. Scale bars represent 1 cm.

In 86 of the 461 families (18%), at least one of the 5 to 7 F_1_ plants showed obvious phenotypic abnormalities (**Fig 2B**), with the most common being chlorosis, necrosis and reduced growth (**Fig 2C**). Such phenotypes are likely to reduce fitness in the field, and are therefore referred to as deleterious. Phenotypic severities ranged from very mild and disappearing with age, to the plant not developing past the cotyledon stage and dying shortly after germination (**Fig 2D**). To test if these phenotypes were heritable, we created pseudo-F_2_ populations for all 86 families in which abnormal phenotypes were observed. To this end, and since *A. arenosa* cannot self-fertilize, the 5 to 7 F_1_ siblings were crossed among each other (sibcrosses) (**Fig 2A**). Most abnormal individuals were not fertile, but those that were, were prioritized for crossing. A total of 37 out of these 86 families tested (43%) produced abnormal offspring that resembled the abnormal parental individuals, indicating that almost half of the initially seen abnormal phenotypes were heritable. The fraction of families with abnormal heritable phenotypes varied considerably between the sampled populations, from 1.7% in the Bridge population to 16.6% in the Steinbruch population (**Fig 2B**, **Table S3**). We did not find a significant correlation between the frequency of abnormal heritable phenotypes and the effective size or inbreeding coefficient of each population (**Table S3**). Taken together, these results show that deleterious abnormal phenotypes segregate at appreciable frequencies in wild *A. arenosa* populations.

### Abnormal *A. arenosa* Plants Show Little Evidence for Autoimmunity

To obtain insights into molecular or physiological processes disturbed in plants with deleterious abnormal phenotypes, we chose two representative families for RNA-seq analysis: A279 from the Sauria population and B772 from the Strečno population (**Fig 2D**, **3A**). In both cases, individuals from these families consistently included plants with stunted growth in either milder or more severe forms, with only the milder forms allowing the formation of a few seeds (**Fig 2D**, **3A, 5A** and **5G**). To allow for comparability between families, tissue was harvested from five normal and five abnormal plants from both families 17 days after germination, when the abnormal phenotype was clearly visible, but before plants expressing the more severe phenotype in the A279 family started to wither (**Fig 3A**). In the A279 family, severely abnormal plants were chosen for expression analysis, while in the B772 family the severity among abnormal plants was not yet evident at the time point we sampled. A PCA showed that most of the variance in gene expression was driven by the difference between plants showing the abnormal phenotype and those that do not (**Fig 3B**, **3C**). This was particularly obvious in the A279 family, where the abnormal phenotype is more pronounced. In abnormal plants of the A279 family, 8,962 genes out of 22,640 annotated genes were differentially expressed, whereas in B772 these were only 1,079 genes (**Fig 3D**). There was substantial overlap (532 genes) between the differentially expressed genes between the A279 and the B772 families (**Fig 3E**). A Gene Ontology (GO) analysis of these overlapping differentially expressed genes (DEGs) showed that these were enriched for the terms “protein phosphorylation” and “response to stress” (**Fig 3F**, **Table S4**). Genes in these two categories were largely upregulated in abnormal plants (**Table S5**). We also assessed the top 100 (**Fig S2A, S2B**) and 500 DEGs for each family separately (**Table S5**). Both sets were enriched for similar terms to those of their intersection, with the A279 DEGs being additionally enriched for “response to abiotic stimulus” and “post-embryonic development”.

**Fig 3.**
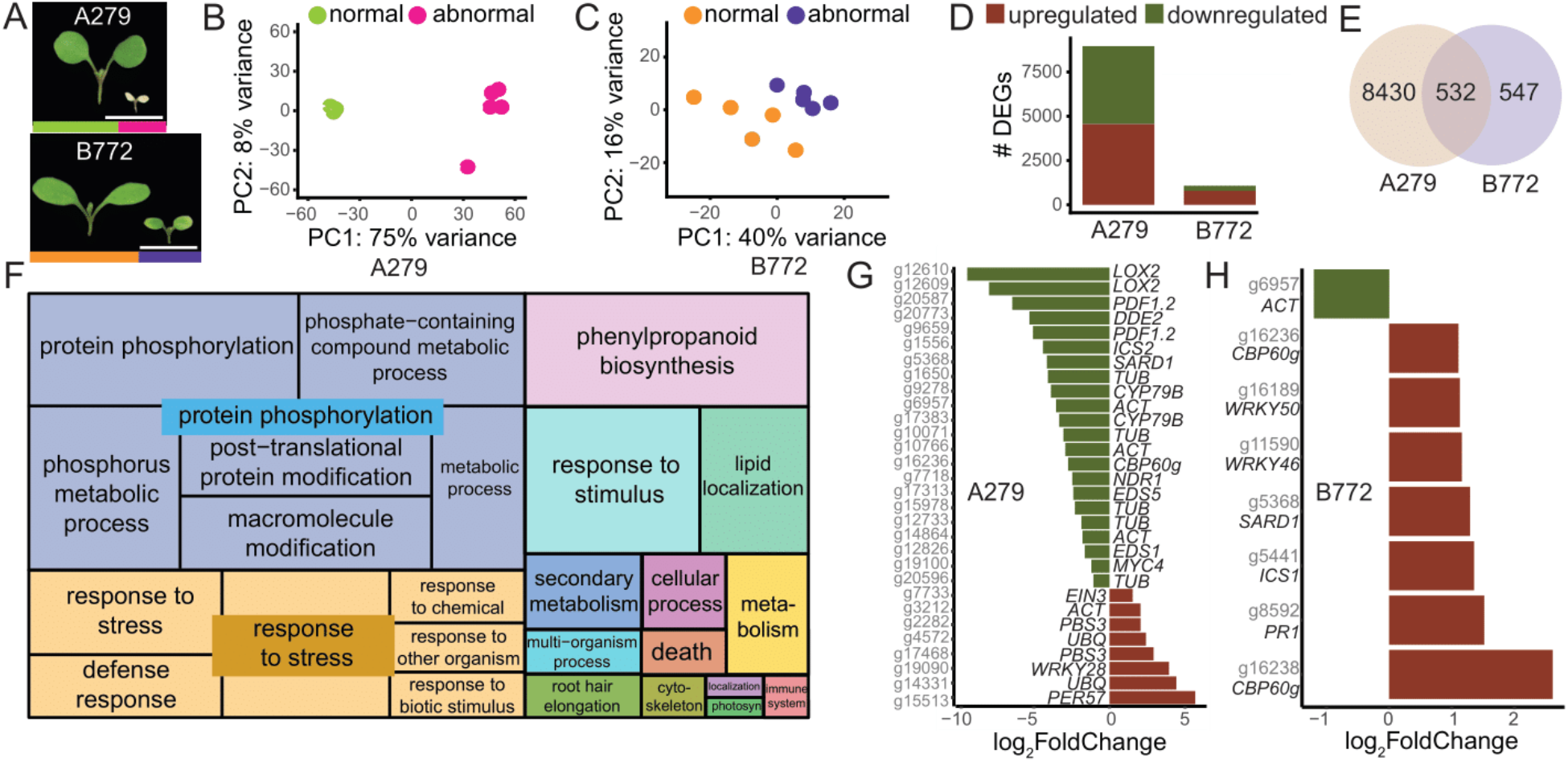
RNA-seq analysis of the A279 and B772 families. **A.** At 17 days after germination, normal (green and yellow) and abnormal (pink and purple) plants from the A279 and B772 families phenotypically differ from each other. Plants were grown at 16°C. Scale bar represents 0.5 cm. **B**, **C.** PCA of gene expression values. The main variance is between normal and abnormal plants. Each dot indicates one biological replicate, with ten (5 normal, 5 abnormal) per family. **D.** Number differentially expressed genes (DEGs) which are either up- or downregulated. **E.** Intersection of DEGs between the A279 and B772 families. **F.** REVIGO Gene Ontology treemap of the DEGs in the intersection between the two families. Size of the square represents -log_10_(*p* value) of each GO term. **G**, **H.** -log_2_FoldChange of significantly (|log_2_FoldChange| >1, padj value < 0.05) DEGs between normal and abnormal plants. *A. thaliana* orthologs of genes chosen for detailed analysis in black. In case of one-to-many ortholog gene associations a representative ortholog or a broader term (e.g. *ACT* for all actin orthologs) is shown.

Our earlier work has shown that defense response genes and genes involved in the salicylic acid (SA) pathway are often upregulated in *A. thaliana* hybrids displaying necrosis and reduced growth^4,25^. When looking at transcriptome-wide differences between incompatible *A. thaliana* hybrids and abnormal *A. arenosa* plants of similar developmental age, we found genes that were differentially expressed in *A. thaliana* hybrids, but not in abnormal *A. arenosa* plants, were enriched for the GO terms “immune response” and “cell death” (**Fig S2C**, **Table S6**). Since these terms did not show up in our unbiased interrogation of differentially expressed genes in *A. arenosa,* we specifically investigated the expression of *A. arenosa* orthologs of such genes, as well as of others, including those related to the synthesis and response to phytohormones^26^ (**Table S7**). Even in the A279 family, with overall many differentially expressed genes, only a few of the 35 selected genes were markedly altered in their expression. Many SA biosynthesis and response genes, including *PR1*, *EDS1*, *ICS1* and *CBP60g*, that were induced in *A. thaliana* hybrids were not upregulated in the abnormal A279 plants (**Fig 3G**). Similarly, only a few pathogenesis related genes were mildly upregulated in abnormal plants in the B772 family (**Fig 3H**). Lastly, since the widespread induction of nucleotide binding site-leucine-rich repeat immune receptor genes (NLRs) was observed in incompatible *A. thaliana* hybrids^25^, the expression of *A. arenosa* NLR orthologs was compared between plants showing an abnormal and a normal phenotype. NLR induction was observed to a far lesser extent than previously seen in *A. thaliana* (**Fig S2D, S2E**, and **Table S8**). In short, individuals with deleterious abnormal phenotypes of both *A. arenosa* families exhibit differential gene expression when compared to their normal siblings. The overall transcriptional profile of these abnormal plants differs from that seen in incompatible *A. thaliana* hybrids, and provides no or only limited evidence for autoimmunity in phenotypically abnormal *A. arenosa* plants. In further support for this, most of the phenotypic abnormalities observed, including those in the A279 and B772 families, were not ameliorated by growing the plants at a higher temperature (**Fig S2F-I**). In contrast, autoimmunity due to hybrid necrosis can often be partially or completely suppressed by growing plants at elevated temperatures^27^.

### A Single Genomic Region is Linked to a Deleterious *A. arenosa* Phenotype

To identify the genetic basis of the deleterious phenotypes observed, both normal and abnormal plants from four independent pseudo-F_2_ families (B772, A279, B635 and B182) were individually genotyped by RAD-seq^21^ (**Fig 2A**). These families were chosen for further examination based on the consistency and severity of their phenotypic defects. Quantitative trait locus (QTL) analysis did not reveal associations between specific genomic regions and the abnormal phenotypes in three out of the four families studied (**Fig S3A-D**).

In the B772 family, however, we identified a single high-confidence QTL on chromosome six. The QTL interval was located between 5.30-5.36 Mb, with a maximum peak at 5.34 Mb and a LOD score of 32.9, containing 12 annotated genes (**Fig 4A, 4B** and **Table S9**). We observed only little recombination, especially among genes found towards the start of the QTL interval in the B772 family (**Fig 4C**). These 12 genes were not significantly (|log_2_FoldChange| >1, padj value < 0.05) differentially expressed between normal and abnormal looking plants (**Table S9**). In a larger genomic region spanning 3.5-8.0 Mb on chromosome 6, four additional marginally significant QTL peaks were found (**Fig 4B**). This larger 4.5 Mb region contained 927 genes (**Table S10**), which were enriched for the GO terms “Cell wall biogenesis” and “Carbohydrate biosynthesis” (**Table S11**). Of these genes, 49 were differentially expressed between normal and abnormal-looking plants (**Table S12**), but these were not enriched for any particular GO term.

**Fig 4.**
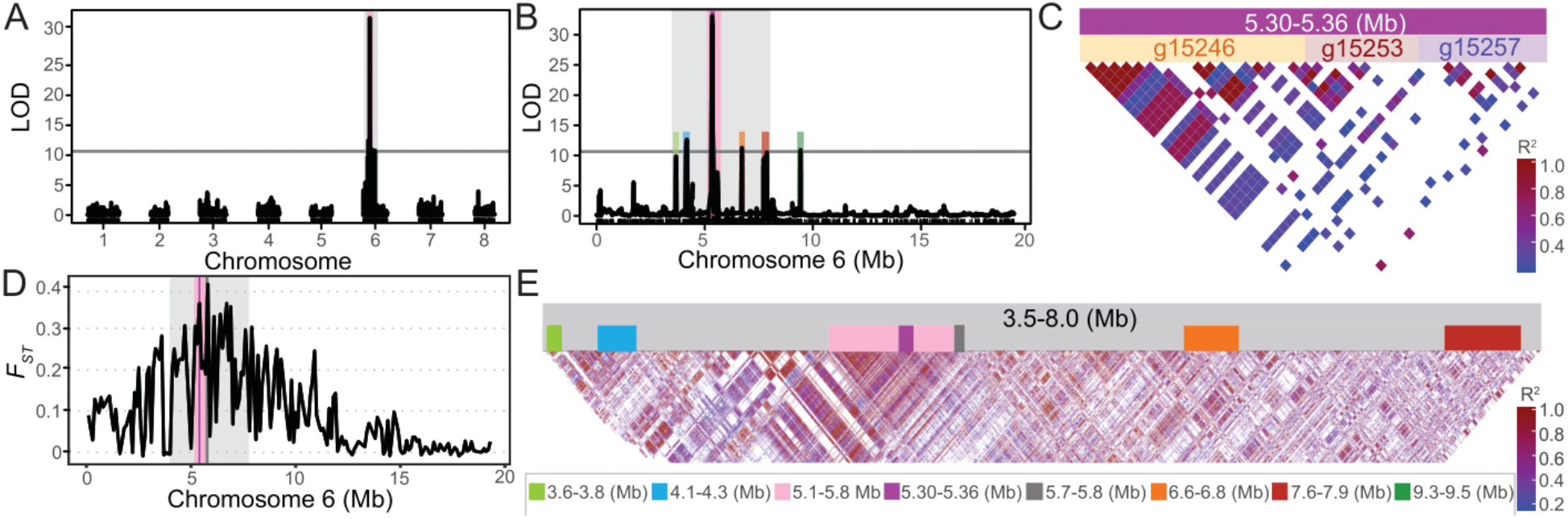
QTL, LD and *F*_*ST*_ analysis of the B772 family. **A**, **B.** A QTL peak is found on chromosome 6 (5.30-5.36 Mb). Horizontal lines indicate 0.05 significance threshold established with 1,000 permutations. The 3.5-8.0 Mb region of chromosome 6 is highlighted in grey. **C.** Linkage disequilibrium (LD) across the 60 kb QTL region between 5.30-5.36 Mb in chromosome 6. Strong linkage is observed in genes found between g15246 and g15253 and, to a lesser extent, in those found until g15257. **D.** Fixation index (*F*_*ST*_) between normal and abnormal plants across chromosome six. The 3.5-8.0 Mb region is highlighted in grey, the 5.30-5.36 Mb region in magenta, the 5.7-5.8 Mb in dark grey and the 5.1-5.8 Mb region in pink. **E.** LD plot for the 3.5-8.0 Mb region in chromosome 6 from all 52 individuals. The region comprising both the highest LOD (magenta) and *F*_*ST*_ (dark grey) is under LD (pink), an indication of reduced recombination. Other colors indicate positions of QTL peaks in B as well as the maximum *F*_*ST*_ value in D.

To examine the genomic region associated with the deleterious phenotype seen in the B772 family more closely, we whole-genome sequenced 52 pseudo-F_2_ individuals, 24 with a normal and 28 with an abnormal phenotype and then calculated the fixation index (*F*_*ST*_) between the two phenotypic groups. Concordant with the QTL analysis described above, a peak in the first half chromosome 6 was observed, with a maximal *F*_*ST*_ value of 0.41 between 5.7-5.8 Mb, followed by 5.3-5.4 Mb with anl *F*_*ST*_ value of 0.36 (**Fig 4D**). These two regions were part of the same 700 kb long linkage block (5.1-5.8 Mb), which showed reduced recombination (**Fig 4E**). No other regions with elevated *F*_*ST*_ values were observed across the entire genome (**Fig S4E**). Taken together, this indicates that a region in the first half of chromosome six, especially a particular LD block, is both genetically differentiated in abnormal plants and linked to their phenotype.

### Some Deleterious *A. arenosa* Phenotypes are Due to Inbreeding Depression

We found a large 700 kb LD block on chromosome 6 to be linked to phenotypic abnormality in the B772 family. The primary cause of these defects may be increased homozygosity in this region, thereby exposing one or multiple deleterious recessive mutations, or incompatibility between two alleles of the same gene^9,10^ or of two closely linked genes.

To distinguish between these possibilities, we investigated the inbreeding coefficient (*F*) within the two phenotypic groups (normal and abnormal) in the 3.5-8.0 Mb region surrounding the LD block on chromosome 6 (**Fig 5A**). Abnormal individuals had a higher *F* than normal ones (**Fig 5B**, **Table S13**). As a control, *F* was calculated for both a similarly sized genomic region on chromosome 1, as well as for the whole genome, and no differences in inbreeding levels between normal and abnormal plants were seen (**Fig S4A, S4B** and **Table S13**).

**Fig 5.**
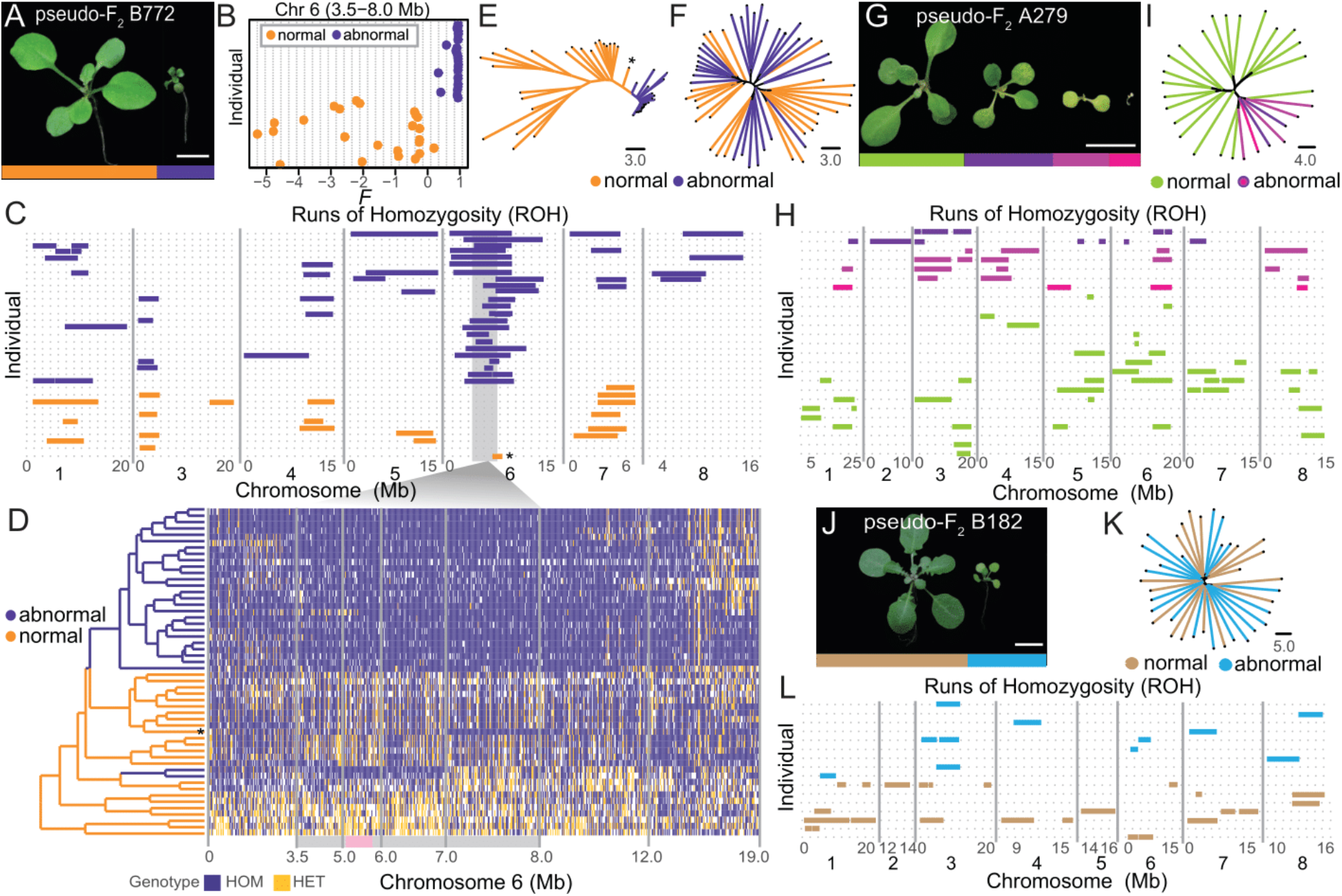
Inbreeding coefficient and runs of homozygosity in the B772, A279 and B182 families. **A.** B772 normal (yellow) and abnormal (purple) plants. **B.** Inbreeding coefficient (*F*) of normal or abnormal plants calculated for the 3.5-8.0 Mb region on chromosome 6. Abnormal plants have a much higher *F*. **C.** Genome-wide runs of homozygosity (ROH). Twenty-three plants with an abnormal phenotype share ROH in chromosome 6, while only one normal looking plant (indicated by an asterisk) had a short ROH in this region. The 3.5-8.0 Mb region is marked in grey. Only plants with at least one ROH are shown. **D.** Genotype calls from normal and abnormal plants in chromosome 6. LD block overlapping the QTL (see Fig 4E) in pink. Homozygous markers (HOM) in purple; heterozygous markers (HET) in yellow. Individuals cluster by chromosome-wide similarity. Dendrogram colors indicate the plant phenotype. Abnormal individuals show higher homozygosity. **E.** NJ tree of the 3.5-8.0 Mb region on chromosome 6. Individuals cluster by phenotype. Branch lengths in nucleotide substitutions per site are indicated. Asterisk indicates the same individual as in C. **F.** NJ tree of the B772 plant’s whole genome. Individuals do not cluster by phenotype. **G.** A279 normal (green), mildly affected (purple), strongly chlorotic (light purple), and severely affected albino (pink) plants. **H.** A279 genome-wide ROH. No ROH is shared across all abnormal individuals. **I.** Genome-wide NJ tree. Individuals cluster by phenotype. **J.** B182 normal (brown) and abnormal (blue) looking plants. **K.** Genome-wide NJ tree. Individuals do not cluster by phenotype. **L.** B182 genome-wide ROH. ROH are not linked to the abnormal phenotype. Plants were between three and five weeks old. Scale bars represent 1 cm.

We also screened for runs of homozygosity (ROH), which would be expected if inbreeding depression would underlie the abnormal B772 phenotype. ROH were identified throughout chromosome 6 in 82-100% of abnormal plants, depending on the parameter settings (**Fig 5C**, **Table S14**). For normal-appearing individuals, only short ROH were identified in this region in one to three (4-12.5%) plants (**Table S14**). For other regions in the genome, ROH were both less frequent and not associated with the occurrence of the abnormal phenotype (**Fig 5C**, **Table S14**). High levels of homozygosity present in abnormal individuals throughout chromosome 6 were confirmed by direct visual inspection of genotype calls (**Fig 5D**). A correlation between homozygosity and the abnormal phenotype was not observed in any other region of the genome (**Fig S4C**). As expected, sequences from the 3.5-8.0 Mb region of chromosome 6, shown to be differentiated between normal and abnormal plants, separated normal and abnormal individuals in a Neighbor-Joining (NJ) tree, which confirms high sequence similarity among abnormal siblings (**Fig 5E**), which was not observed for the whole genome (**Fig 5F**), or for the control region on chromosome 1 (**Fig S4C**).

To identify candidate genes underlying the abnormal phenotype, we screened for high-impact mutations. We found seven such mutations in the 3.5-8.0 Mb region of chromosome 6, six of which were homozygous, although not exclusively, in abnormal plants (**Table S15**). This indicates that a mutation in more than one gene in this region may be underlying the phenotypic defects, or that mutations other than those defined as having high impact are relevant.

For the families where we did not find a clear QTL associated with abnormal siblings, we wondered whether simultaneous homozygosity in multiple regions was causing the abnormal phenotypes, possibly explaining why no QTLs were identified before. To investigate this, we sequenced the complete genomes of individuals from the A279 family, which showed a range of different defects. Of the 28 sequenced individuals, two were mildly chlorotic and slightly smaller in size, four were strongly chlorotic, dwarfed and did not develop true leaves, and one was a very small albino (**Fig 5G**). We calculated *F* in 5 Mb windows across the whole genome, and saw that there are multiple regions where abnormal plants tended to show a higher *F* than normal plants (**Fig S4D**, **Table S16**). We also calculated genome-wide *F*, and abnormal plants were part of the upper quartile of values (**Fig S4E**, **Table S16**). Different from the B772 family, we found ROH in similar proportions in both normal and abnormal plants (**Fig 5H, Table 17**). There was no single ROH that was shared between all abnormal A279 individuals however, which was also apparent with direct visual inspection of genotype calls (**Fig S4G**). The data is consistent with different combinations of unlinked ROH giving rise to different phenotypic defects, but more individuals would need to be analyzed to more fully support this assertion. Unlike B772, abnormal plants cluster together when whole-genome sequence variation is considered (**Fig 5I**), further hinting at multiple genomic stretches contributing to the phenotypic abnormalities.

Finally, we whole-genome sequenced a third family, B182 (17 normal, 20 abnormal plants) from the Horse population, which consistently segregated plants with a chlorotic dwarf phenotype, which did not vary in severity (**Fig 5J**), but in which we also had failed to detect a clear QTL for the abnormal phenotype. Unlike the two previous families, no relationship between the abnormal phenotype and overall inbreeding levels, or ROH were found, nor were the abnormal plants genetically more similar to each other than to normal individuals (**Fig 5J-L**, **S4H** and **S4I**, **TableS18** and **S19**). We therefore suspect a more complex genetic architecture underlying the phenotypic segregation in the B182 family, which is supported by many more than a quarter of plants, as expected for a single causal genomic region, having an abnormal phenotype. To summarize, we found high homozygosity present in particular genomic regions of abnormal plants, to be linked to their deleterious phenotype.

### Homozygosity in a Region Underlying Phenotypic Abnormalities is Rare in the Wild

Genomic areas under positive selection can be identified by scanning for loci that show an excess proportion of identity by descent among individuals in a population^28,29^. The same principle could be used to scan for regions under negative selection, because they should be rarer in a population, and would be candidate loci for acting as sources of inbreeding depression. We looked at homozygosity in wild individuals from the Strečno population, where the original B772 mother plant was collected from, and asked how often ROH were observed across chromosome 6, especially in the 3.5-8.0 Mb region associated with the presence of the abnormal phenotype. We whole-genome sequenced 40 individuals (**Fig 1A**, **Table S1**) and saw that most *F* values ranged from -0.3 to 0.2 per individual (**Fig 6A**), slightly lower than what we had inferred from our RAD-seq data (**Fig 1D**, **Table S20**). Genome-wide *F* was not unusually high in the original B772 F_0_ individual collected from the wild, compared to other Strečno individuals. The 3.5-8.0 Mb region of chromosome 6, which we found to be homozygous in abnormal pseudo-F_2_ B772 plants, was not exceptionally homozygous in the B772 plant either (**Fig 6B**). This agrees with our observation of the B772 F_0_ plant appearing normal in the field. Four individuals showed positive *F* values in this 3.5-8.0 Mb region, and this was not due to missing data, which was below 0.02% across chromosome 6 for all individuals (**Table S20**). Among these four, the most homozygous individual was B771, reaching a *F* value of almost 1 (**Fig 6B**, **Table S20**), and which had been found right next to the B772 individual. Extensive homozygosity in the B771 plant across chromosome 6 was confirmed by directly inspecting genotype calls (**Fig 6C**). As a control, *F* was calculated for the 3.5-8.0 Mb region in chromosome 1 across all 40 plants (**Fig 6D**). Neither B772 nor B771 had high *F* in this region, but three other plants had, indicating that homozygous stretches across the genome are relatively common in wild individuals from this population, an observation confirmed by visually inspecting genotype calls (**Fig S5A**). A genome-wide NJ tree, as well as a PCA, placed B771 and B772 in close proximity to each other (**Fig 6E, 6F**). These analyses revealed three distinct genetic groups present in the Strečno population (**Fig 6E, 6F**, **Table S20**), which is in line with largely positive genome-wide Tajima’s D values (**Fig S5B**), indicating, *en gros*, the presence of multiple alleles found in the Strečno population at intermediate frequencies.

**Fig 6.**
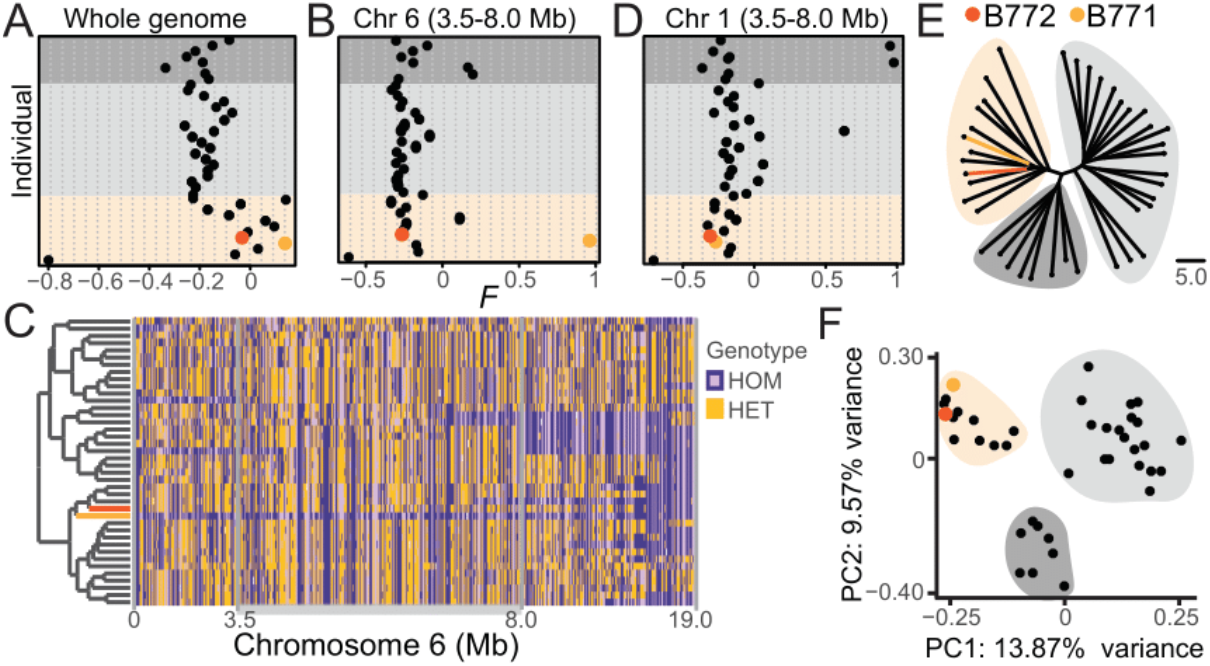
Homozygosity in wild Strečno individuals. **A**, **B** and **D.** Inbreeding coefficient (*F*) per plant for the whole genome (A), the 3.5-8.0 Mb region of chromosome 6 (B), and the 3.5-8.0 Mb region of chromosome 1 (D), a control region. Individuals follow the same order in the three plots. B771 (yellow) but not B772 (orange) had a high *F* in the 3.5-8.0 Mb region of chromosome 6. Background colors as in E. **C.** Chromosome 6 genotype calls from the 40 plants. Homozygous markers (HOM) in purple; heterozygous markers (HET) in yellow. Individuals cluster by similarity. B771 but not B772 shows high homozygosity. The 3.5-8.0 Mb region is in grey. **E.** Genome-wide NJ tree. Three genetic groups are visible (dark grey, light grey and yellow), with B771 and B772 belonging to the same group (Table S20). Branch lengths in nucleotide substitutions are indicated. **F.** Genome-wide PCA, each individual is represented by a dot. Clustering of individuals and colors as in E.

In brief, we found homozygosity in the 3.5-8.0 Mb region of chromosome 6 to be rare in wild plants, in agreement with it imparting a fitness penalty. Homozygous stretches in other regions of the genome, however, were common in the wild.

## Discussion

### Balancing Inbreeding and Outbreeding Depression

The genetic distance between any two individuals can negatively impact their offspring’s fitness^30^. On one end, high divergence between individuals may lead to allelic mismatches and outbreeding depression. Such deleterious epistatic genetic interactions are likely a consequence of breaking up co-evolved gene complexes^31^. On the other end, the progeny of parents that are genetically relatively similar to each other can exhibit signs of inbreeding depression, which occurs when recessive deleterious mutations become visible when in a homozygous state, or when heterozygote advantage is lost^14,20^. These two opposing phenomena - outbreeding and inbreeding depression - indicate that an optimum parental genetic distance may exist^32,33^. Indeed, a study of human fertility in a large population with completely known pedigrees has suggested that there is an optimal degree of relatedness for fitness^34^. This was also shown in model organisms for three major eukaryotic groups, including *A. thaliana*, where the expected hump-shaped relationship between genetic distance and fitness was observed^35^.

### Likelihood of Inbreeding or Outbreeding Depression Occurring

A key factor for whether an organism is more prone to experience inbreeding or outbreeding depression, is its mating system. In selfing organisms, effective purging of deleterious alleles reduces the likelihood of inbreeding depression and high genetic drift can enhance differentiation between populations, thereby increasing the chances of outbreeding depression^2^. In contrast, outcrossing species, which are often less affected by genetic drift, can tolerate a higher load of deleterious alleles, as long as these are rare and do not reduce fitness as heterozygotes, making inbreeding depression more likely to become problematic^14^.

A second factor likely to influence whether plants are more or less likely to suffer from inbreeding or outbreeding depression is ploidy. In outcrossers, tetraploid plants can have up to twice the number of alleles than in diploid plants, potentially providing much more opportunity for deleterious interactions between divergent alleles to occur, resulting in outbreeding depression. In addition, polyploidy increases heterozygosity, thereby masking recessive deleterious alleles or maintaining overdominance^36–38^, reducing the likelihood of inbreeding depression. Because both diploid and tetraploid cytotypes exist, *A. arenosa* would be a good system for studying whether outbreeding depression is indeed more common than inbreeding depression in polyploids.

### Rare Occurrence of Hybrid Necrosis in *A. arenosa*

In the *A. arenosa* congener *A. thaliana*, hybrid necrosis is by far the most common F_1_ hybrid weakness phenotype observed under laboratory conditions, usually resulting from crosses between individuals from different populations^4^, but other incompatibilities have been observed as well, including single-locus F_1_^39^, two-locus F_1_^40^, or two-locus F_2_ incompatibilities^41–45^. We have found the incidence of abnormal phenotypes in *A. arenosa* material collected from the wild, and raised in the lab, to be relatively high. While the spectrum of abnormal phenotypes in our material is similar to that of *A. thaliana*^4,9^, albeit at higher frequencies, hybrid necrosis may likely not be the most common genetic cause for these phenotypic abnormalities, but rather inbreeding depression. We note however, that we have focused on naturally occurring crosses within local populations. Since breaking up co-evolved loci can result in hybrid necrosis, incompatibility between non-occurring individuals, such as those originating from different populations, may be more common. We have performed a limited number of crosses (n=33) between *A. arenosa* individuals from different populations. Here, abnormal phenotypes were even more common, with half of all crosses segregating abnormal phenotypes (**Table S21**). Further study will reveal whether these phenotypes are linked to hybrid necrosis.

### Phenotypic Abnormalities as a Result of Inbreeding Depression

Inbreeding depression commonly occurs in outcrossing plant species^46^, and at least some of the deleterious *A. arenosa* phenotypes we observed can likely be attributed to inbreeding depression, because they are linked to extended homozygosity. These abnormal and heritable phenotypes were common, and observed in about one in ten maternal sibships. In a study of another wild, outcrossing Brassicaceae species, *Leavenworthia alabamica*, a third of all families were lost after one generation of selfing^47^, highlighting the strong impact inbreeding can have in plant fitness. This study reported around 3% of abnormal phenotypes among all individuals after outcrossing. Of 2,640 *A. arenosa* plants raised from seeds collected in the wild, we observed morphological abnormalities in 157 plants (5.9%), which is higher, but not hugely disparate. A related question is that of hybrid vigor, or heterosis^14,48^, and how common it is in crosses within populations compared to crosses between populations. If our observations reflect a high degree of inbreeding depression in the studied *A. arenosa* populations, we expect heterosis to be more frequent in crosses between populations, since recessive deleterious mutations will be masked, or heterozygote advantage gained.

### Different Genetic Architectures Underlie Phenotypic Abnormalities in *A. arenosa*

Of the families we investigated more in detail, relatively simple genetics was only found in the case of the B772 family, where a single clear QTL was part of a larger LD block, suggesting reduced recombination in this genomic region, which is known to lead to the increased accumulation of deleterious mutations^49^. This specific QTL region was found to be homozygous in all abnormal plants, but largely heterozygous in all normal plants. Whether the phenotypic abnormalities in the B772 family are due to deleterious mutations found in one or multiple genes in the QTL interval is unclear. In the causal region, we found high-impact mutations in several genes, but none of these were perfectly correlated with the phenotypic abnormalities.

Homozygous genomic stretches were found to be relatively common in wild individuals. The degree of observed homozygosity is not entirely unexpected, since fertilization typically occurs between nearby individuals, which are more likely to share a recent common ancestor. In the specific case of the B772 family collected from Strečno, we found that the B772 F_0_ individual collected in the wild was not homozygous for the region underlying the abnormal phenotype, but the B771 F_0_ individual growing right next to it was. It is conceivable that many of the B772 offspring result from fertilization by B771 pollen. That out of 40 plants only the B771 plant showed ROH in the 3.5-8.0 Mb region of chromosome 6 supports our observation of homozygosity here being disadvantageous. We were able to collect only very few seeds (≤10) from this plant, perhaps another indication of reduced fitness brought on by ROH in chromosome 6. We note however, that B771 had a high genome-wide *F*, so we cannot exclude that other regions in the genome may have affected its ability to produce seeds.

In the other families studied in detail, QTL analysis did not identify regions of the genome that were clearly linked to the abnormal phenotype. The most parsimonious explanation is the involvement of multiple genomic regions, with different combinations of homozygous regions leading to the different phenotypic abnormalities observed. This coincides with previous observations that small-effect mutations distributed across multiple loci, which are maintained in natural populations at low frequencies, collectively can make for a common source of inbreeding depression^14^. Perhaps the most intriguing case is that of the B182 family, where no obvious QTL or ROH linked to the phenotype was detected. One explanation could be that the family is fixed for one or more causal genes, but that the phenotype has only limited penetrance, i.e., shows up only in a fraction of individuals, or that the expression of the phenotype is epigenetically unstable.

### Reducing Inbreeding Depression Through Conservation Genomics

Accelerated climate and land use change are prominent drivers of increased population fragmentation in many species^50^. Inbreeding depression in turn tends to be especially problematic in small, fragmented populations^51^. The wild *A. arenosa* populations in this study were neither particularly small nor fragmented (**Table S1** and **S3**), yet the negative consequences of inbreeding were clearly visible. Future studies with species that experience more or less population fragmentation than *A. arenosa* will inform us on whether and how the extent of population fragmentation and population differentiation affect the genetic architecture of inbreeding depression. Such knowledge can help optimize conservation strategies such as genetic rescue, which will be much simpler if deleterious phenotypes are due to a small number of genes or genomic regions^52–54^, in line with genomics becoming a central pillar of conservation management^55^. Finally, regardless of underlying ecology, our work has shown that it can be highly profitable to closely study the entire range of phenotypes in individuals collected from the wild and raised in a common garden.

## Methods

### Plant material

Plants were stratified in the dark at 4°C for 5-8 days prior to planting on soil. Plants were grown in long days (16 h of light) at 16°C or 23°C at 65% relative humidity under 110 to 140 μmol m^−2^ s^−1^ light provided by Philips GreenPower TLED modules (Philips Lighting GmbH, Hamburg, Germany) for six to eight weeks and were later moved to greenhouse conditions. The ploidy of each population sampled was estimated via flow-cytometry (one representative plant per population) and using nQuire^56^. Plants in the wild were sampled under permit number (062-219/18).

### De novo genome assembly and annotation

A single *A. arenosa* plant (701a) from the Strečno population was grown as described above. Fresh green tissue was harvested over several weeks. High molecular weight DNA was extracted from ~50 ml finely ground tissue powder: nuclei were enriched by gentle resuspension of tissue powder in 500 ml fresh and ice-cold isolation buffer (10mM Tris pH8, 0.1M KCl, 10mM EDTA, 0.5M sucrose, 4mM spermidine, 1mM spermine), followed by filtration through two layers of miracloth, gentle addition of 25 ml isolation buffer containing 20% Triton-X-100, incubation on ice for 15 min, and centrifugation at 3,000 g. Nuclei pellets were washed with isolation buffer containing 1% Triton-X-100, gently resuspended in 30 ml G2 lysis buffer (Qiagen), and incubated with 50ug/ml RNaseA (Qiagen) at 37°C for 30 min, followed by proteinase K treatment (200ug/ml; Qiagen) at 50°C overnight. After centrifugation at 8,000 g, DNA-containing supernatant was purified with Qiagen genomic tip 100 following the manufacturer’s instructions. 0.7 volumes of isopropanol were gently added to the flow-through, and DNA spooled with slow tube rotations, then resuspended in EB buffer (Qiagen) at 4°C. A 20-30 kb XL library was sequenced on a PacBio RS II instrument (Pacific Biosciences, Menlo Park, CA, USA). A second >20 kb library (from the same genomic DNA) was sequenced on a PacBio Sequel instrument with Binding Kit 3.0. In addition, a PCR-free library was prepared with the NxSeq® AmpFREE Low DNA Library Kit (Lucigen, Middleton, WI, USA) according to the manufacturer’s instructions, and sequenced on an Illumina HiSeq3000 instrument (Illumina, San Diego, USA) in paired-end mode (2x 150 bp). PacBio long-reads were assembled with Falcon (v0.3.0)^57^. The resulting contigs were polished using the PCR-free reads with Quiver/Arrow (v2.3.2; https://github.com/PacificBiosciences/GenomicConsensus), as well as with Pilon (v1.16)^58^. Duplicate contigs were then removed using HaploMerger2 (v3.4)^59^. The polished contigs were then scaffolded based on the *A. lyrata* assembly (v1)^60^ using REVEAL (v0.2.1)^61^. *A. arenosa* transcriptome sequencing data were mapped against the scaffolded genome assembly using HISAT (v2.0.5)^62^. Subsequently, the mapping results were used as extrinsic RNA sequencing evidence when annotating the genome using AUGUSTUS (v3.3.3)^63^. Transposable elements and repetitive regions were identified using RepeatModeler2 (v2.01)^64^ and subsequently masked using RepeatMasker (v4.4.0; http://repeatmasker.org). Orthologous genes shared between *A. arenosa* and the *A. thaliana* reference annotation (Araport11) were identified using Orthofinder (v2.4.0)^65^. *A. arenosa* and *A. thaliana* protein fasta files were subsetted to only keep the primary transcript for orthologous assignment using the AGAT toolkit (v0.4.0) (https://github.com/NBISweden/AGAT).

### Genotyping-by-sequencing and QTL mapping

Genomic DNA was extracted from plants with CTAB (cetyl trimethyl ammonium bromide) buffer^66^ and then purified through chloroform extraction and isopropanol precipitation^67^. RAD-seq Genotyping-by-sequencing (GBS) was performed using KpnI tags^21^. Briefly, libraries were single-end sequenced (1×150 bp) on an Illumina HiSeq3000 instrument. Reads were processed with Stacks (v1.35)^68^ and mapped to our *A. arenosa* reference (**Table S2**) with bwa-mem (v0.7.15)^69^, variant calling was performed with GATK (v4.0)70 and SNP filtering with VCFtools (v0.1.14)^71^. Filtering criteria for F_0_ individuals were to retain bi-allelic SNPs only, SNPs with at most 10% missing data, individuals with less than 40% missing data, and SNPs with a minimum depth of 4 and a maximum depth of 80. SNPs were then pruned (--indep-pairwise 50 5 0.5) with PLINK (v1.90)^72^. Downstream analyses were based on 345 individuals and 5,199 markers with an average depth per position of 45 (**Fig 1** and **S1**). For pseudo-F_2_ individuals, SNPs with 30% missing data, individuals with less than 40% missing data and SNPs with a minimum allele frequency of 0.01 were removed, and so were those with a depth of more than 100. Pseudo-F_2_ plants were used as mapping populations for QTL analyses, which were performed with R/qtl^73^. Here, the genome was scanned with a single QTL mode (scanone) using the expectation–maximization (EM) algorithm. QTL analyses were based on 227 individuals and 11,858 markers with an average depth per position of 29 (B772 family), 162 individuals and 9,064 markers with an average depth per position of 23 (B182 family), 183 individuals and 14,672 markers with an average depth per position of 25 (A279 family) and 271 individuals and 6,110 markers with an average depth per position of 25x (B635 family) (**Fig 4** and **S3**).

### RNA sequencing

Five biological replicates of 21 day-old (17 days after germination) shoots of normal and abnormal B772 and A279 plants were collected. After RNA extraction^74^, sequencing libraries were prepared using the TruSeq Total RNA Kit (Illumina) and the Ribo-Zero Plant Kit (Illumina). Libraries were paired-end sequenced on an Illumina HiSeq3000 instrument. Reads were mapped against our *A. arenosa* reference genome using bowtie2 (v2.2.6)^75^. Default parameters were chosen unless mentioned otherwise. Transcript abundance was calculated with RSEM (v1.2.31)^76^. Differential gene expression analyses were performed using DESeq2 (v1.18.1)^77^. Genes with less than ten counts over all samples were removed from downstream analyses. Significant changes in gene expression between normal and abnormal plants from the same family were determined by filtering for genes with a |log_2_FoldChange| >1 and padj value < 0.05. Plots were generated using the R package ggplot2 (v3.2.0)^78^. Gene Ontology (GO) analyses were performed using AgriGO^79^ with the SEA method (default settings), using TAIR9 as a gene model and taking as input the *A. thaliana* orthologs of the annotated *A. arenosa* genes. GO results were visualized with REVIGO treemap^80^, for clearer visualization a maximum of 15 GO terms were plotted. The complete list of GO terms is found in Table S4 and S5. Transcriptome-wide expression changes between incompatible *A. thaliana* hybrids and their parents^25^ were compared to expression changes between normal and abnormal *A. arenosa* individuals through gene orthology assignment. For every differentially expressed gene in either the *A. arenosa* or *A. thaliana* dataset, the corresponding orthogroup was extracted. Intersections were then created between orthogroups containing differentially expressed genes (**Fig S2C, Table S6**).

### Whole-genome sequencing

Libraries from 52 pseudo-F_2_ individuals (23 normal, 29 abnormal) from the B772 family, 28 pseudo-F_2_ individuals (7 abnormal, 19 normal) from the A279 family, 37 pseudo-F_2_ individuals (17 normal, 20 abnormal) from the B182 family, and 40 wild F_0_ individuals from the Strečno population were prepared ^81^, and were paired-end sequenced on an Illumina HiSeq3000 instrument. Reads were processed with Stacks (v1.35)^68^ and mapped to our *A. arenosa* reference genome with bwa-mem (v0.7.15)^69^, variant calling was performed with GATK (v4)^70^. SNPs with at most 30% missing data (50% for the A279 family), individuals with less than 35% missing data, SNPs with a minimum allele frequency of 0.01, a minimum depth of 4 and a maximum depth of 100 were filtered out, resulting in 43,885 SNPs (B772 pseudo-F_2_), 138,620 SNPs (A279 pseudo-F_2_), 45,550 SNPs (B182 pseudo-F_2_) and 57,922 SNPs (Strečno F_0_). After pruning (--indep-pairwise 50 5 0.5) 3,737 SNPs with an average depth per position of 16 (B772 pseudo-F_2_), 7,037 SNPs with an average depth per position of 18 (A279 pseudo-F_2_), 4,987 SNPs with an average depth per position of 13 (B182 pseudo-F_2_) and 3,903 SNPs with an average depth per position of 10 (Strečno F_0_) remained.

### Population genetic analyses

Principal component analyses were calculated with smartPCA^82^. *F*_*ST*_ and Tajima’s D were determined with VCFtools (v0.1.14). Maps were created with the R-package ggmap (v3.0)^83^. Ancestral populations were estimated using ADMIXTURE^24^. Effective population sizes were estimated with NeEstimator (v2)^84^, using the Linkage disequilibrium model. Linkage disequilibrium (r^2^), inbreeding coefficient (*F*) and runs of homozygosity (ROH) were calculated with PLINK (v1.90). For ROH identification, both default parameters (--homozyg-window-het=1 and --homozyg-window-missing=5, **Fig 5C**, **Table S14**) as well as slightly less stringent filters (--homozyg-window-het=2 and --homozyg-window-missing=15, **Fig 5H, 5L**, **Table S17** and **S19**) were used. Sequences were aligned with MUSCLE (v3.8.31)^85^ and then visualized with Aliview (v1.18.1)^86^. Neighbor-Joining trees were estimated with either Jalview^87^ or Fastphylo (v1.0.1)^88^, and visualized in Figtree (v1.4.3)^89^. Genotypes from VCF files were visualized with Genotype Plot (https://github.com/JimWhiting91/genotype_plot). SnpEff^90^ was used to predict the effects of variants and identify high impact mutations using our *A. arenosa* annotation as a reference.

## Data Availability

Sequencing data can be found at the European Nucleotide Archive (ENA) under project numbers PRJEB42608 (RNA-seq experiment) and PRJEB42625 (*A. arenosa* assembly and annotation).

## Acknowledgements

We thank Christa Lanz, Josip Perković, Hung Vo and Frank Vogt for technical support, Wei Yuan for guidance with the RNA extractions, and Gautam Shirsekar and Sergio Latorre for discussion. We also thank Patrick Hüther and Claude Becker for their help with sample transportation logistics.

## Author contributions

Conceptualization: ACB, FK, DW. Formal analysis: ACB, MC. Funding acquisition: FK, DW. Investigation: ACB, MC, RS, MK, IB, FB, DP, FK. Project administration: DW. Supervision: DW. Writing – original draft: ACB. Writing – review & editing: ACB and DW with contributions from all authors.

## Supplemental Material

### Supplemental Tables

All Supplemental Tables are found in a separate. xlsx file.

**Fig S1.**
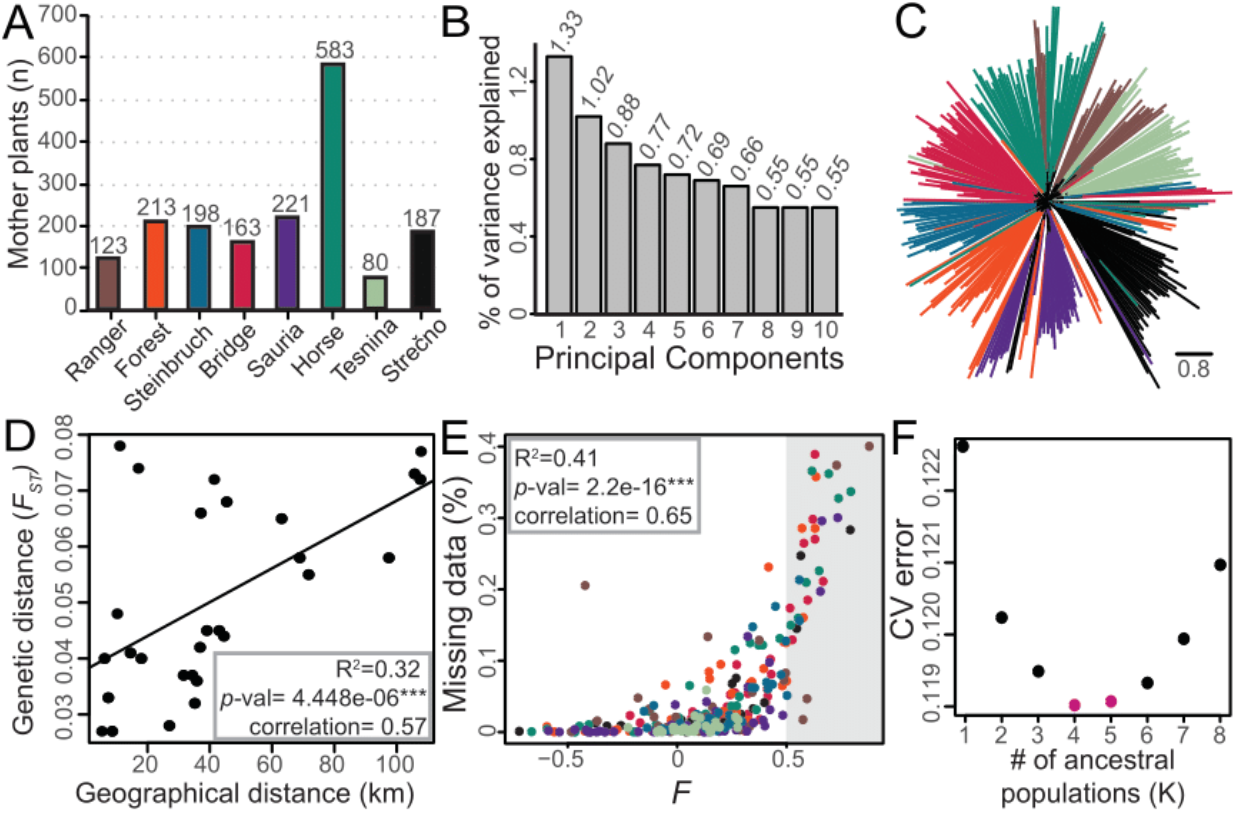
Information on sampled *A. arenosa* populations. **A.** Number of plants sampled in each population. **B.** Principal components and the percentage of variance explained by each. Related to Fig 1B. **C.** Neighbour-Joining (NJ) tree of the 345 sequenced individuals, colored by population. Colors as in A. **D.** Pearson Correlation Coefficient between genetic (*F*_*ST*_) and geographical distance among populations. **E.** Correlation between the amount of missing data per individual and *F*. *F* values above 0.5 are highlighted in grey. Colors as in A. **F.** ADMIXTURE cross-validation error. K=4 and K=5 have the lowest errors (pink). Related to Fig 1.

**Fig S2.**
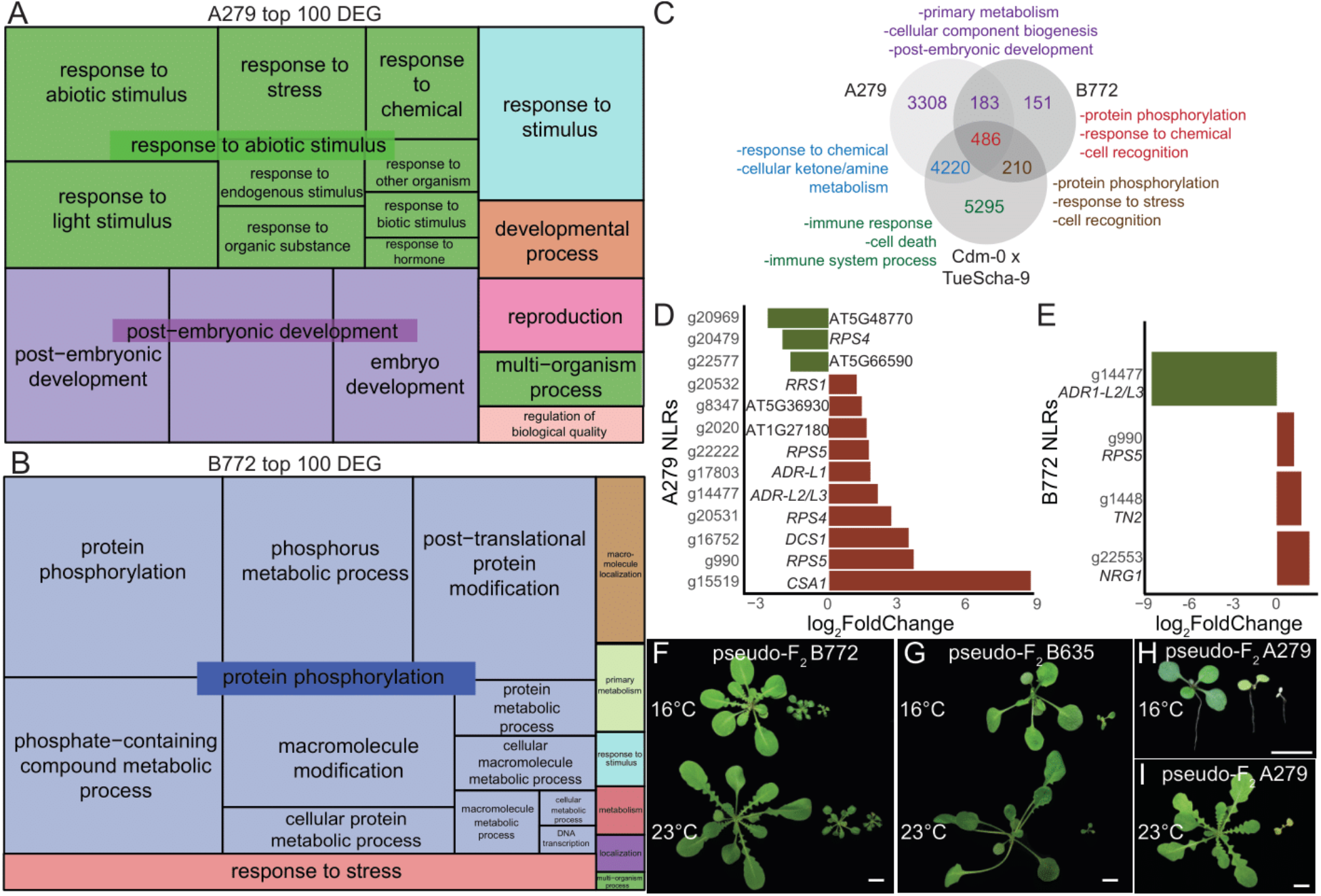
RNA-seq analysis of the A279 and B772 families and pseudo-F_2_ phenotypes. **A**, **B.** REVIGO Gene Ontology treemap of the top 100 DEGs between normal and abnormal plants in the A279 (A) and B772 (B) families. Size of the square represents -log_10_(*p* value) of each GO term. **C.** Intersection of DEGs between the A279 and B772 *A. arenosa* families and the incompatible Cdm-0 x TueScha-9 F_1_ *A. thaliana* hybrid ^25^. Some DEGs are exclusively present in *A. thaliana* (green), *A. arenosa* (purple), and some are found in both (red). The top non-redundant GO categories for each intersection are shown. **D**, **E.** Significant (|log_2_FoldChange| >1, padj value < 0.05) NLR ortholog expression changes between normal and abnormal plants in A279 (D) and B772 (E). *A. thaliana* orthologs in black. In case of one-to-many ortholog gene associations a representative ortholog is shown (Table S8). **F**-**I.** Pseudo-F_2_ plants from the B772 (F), the B635 (G), and the A279 (H, I) families. Plants were either grown for four weeks at 16°C and then transferred to 23°C for another week (F, G, I), or grown at 16°C for four weeks (H). Temperature had no apparent effect on the severity of the abnormal deleterious phenotypes in *A arenosa*. Scale bar represents 1cm. Related to Fig 3.

**Fig S3.**
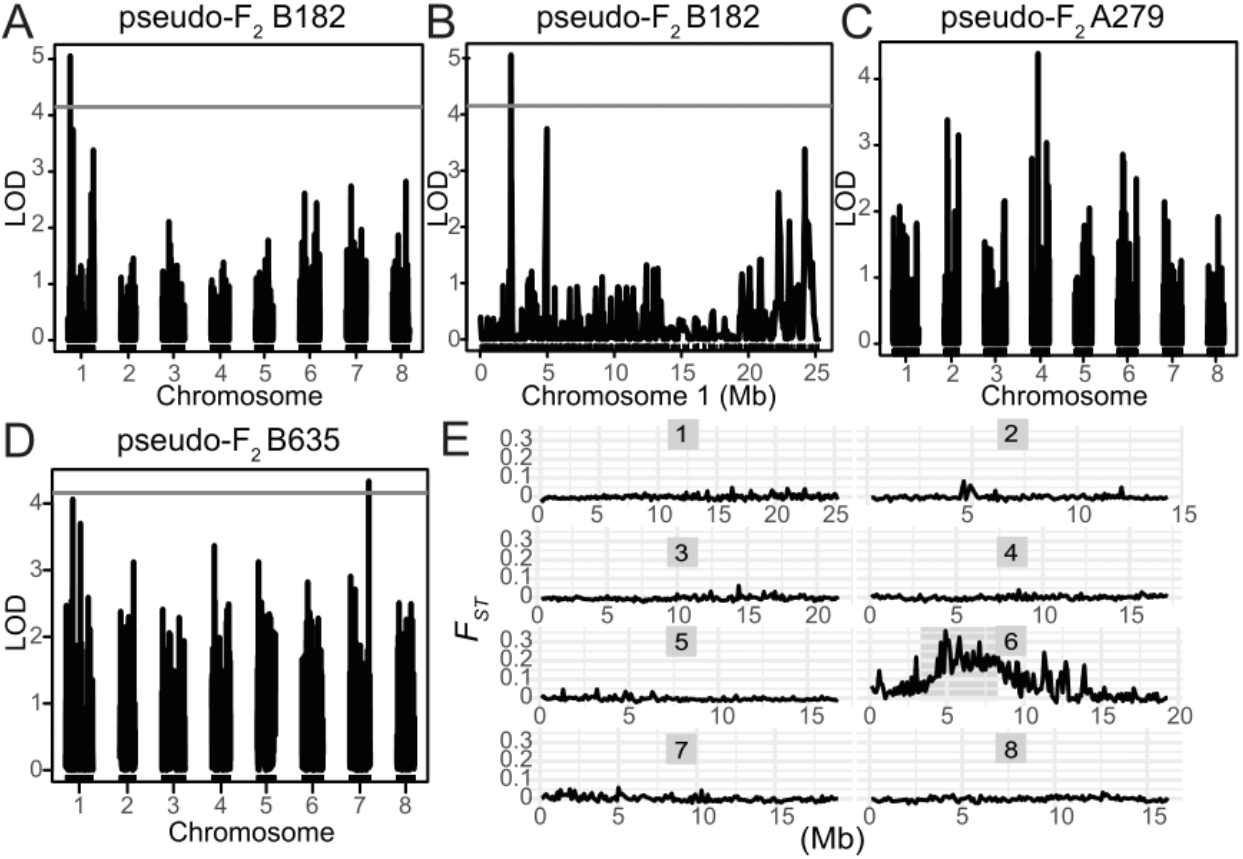
Pseudo-F_2_ QTL analyses and genome-wide *F*_*ST*_. **A**-**D.** QTL analyses for the B182 (A, B), A279 (C) and B635 (D) families. Horizontal lines indicate 0.05 significance threshold established with 1,000 permutations. The absence of this line indicates the significance threshold is above the plotted values. No clear QTLs were found. **E.** Genome-wide fixation index (*F*_*ST*_) between 24 normal and 28 abnormal plants from the B772 family. Higher values are seen exclusively in the first half of chromosome six. The 3.5-8.0 region in chromosome 6 is highlighted in grey. Related to Fig 4.

**Fig S4.**
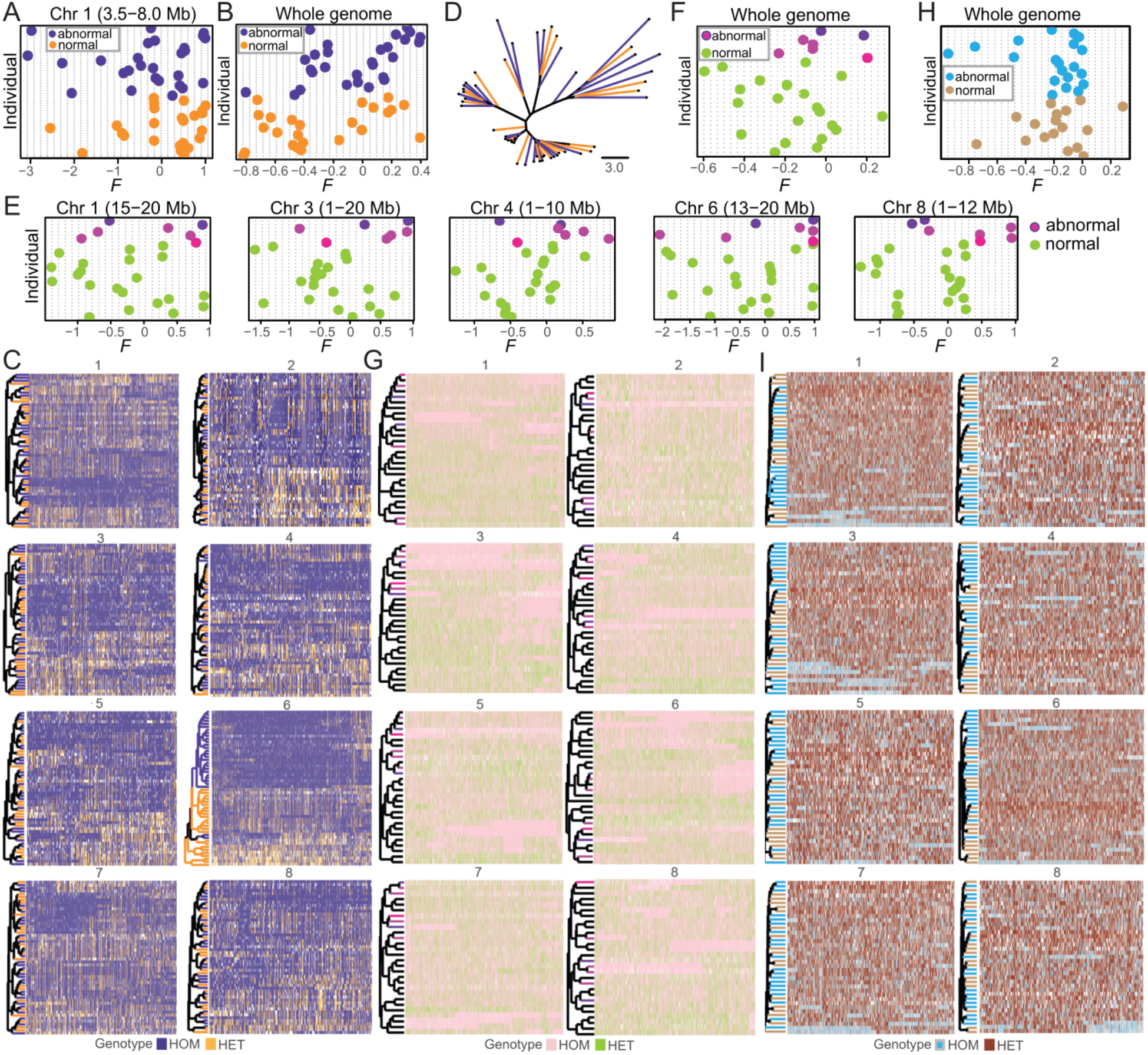
Inbreeding coefficient and ROH in the B772, A279 and B182 families. **A**, **B.** Inbreeding coefficient (*F*) of B772 plants showing an abnormal (purple) and normal (yellow) phenotype calculated for a control region, 3.5-8.0 Mb on chromosome 1 (A), and for the whole genome (B). **C**, **G** and **I.** Genome-wide genotype calls from normal and abnormal B772 (C), A279 (G) and B182 (I) plants. Each chromosome is plotted independently. Individuals cluster by chromosome-wide similarity. Dendrogram colors indicate the plant phenotype (see Fig 5). **D.** NJ tree for the 3.5-8.0 Mb region of chromosome 1 in B772 plants. Individuals do not cluster by phenotype. Branch lengths in nucleotide substitutions per site are indicated. **E.** Genomic regions where *F* tends to be higher in abnormal A279 plants. **F** and **H.** Genome-wide *F* in A279 (F) and B182 (H) plants. Abnormal A279 plants are part of the upper quartile of *F* values across all 28 plants, while in abnormal B182 plants do not tend to have a higher *F.* Related to Fig 5.

**Fig S5.**
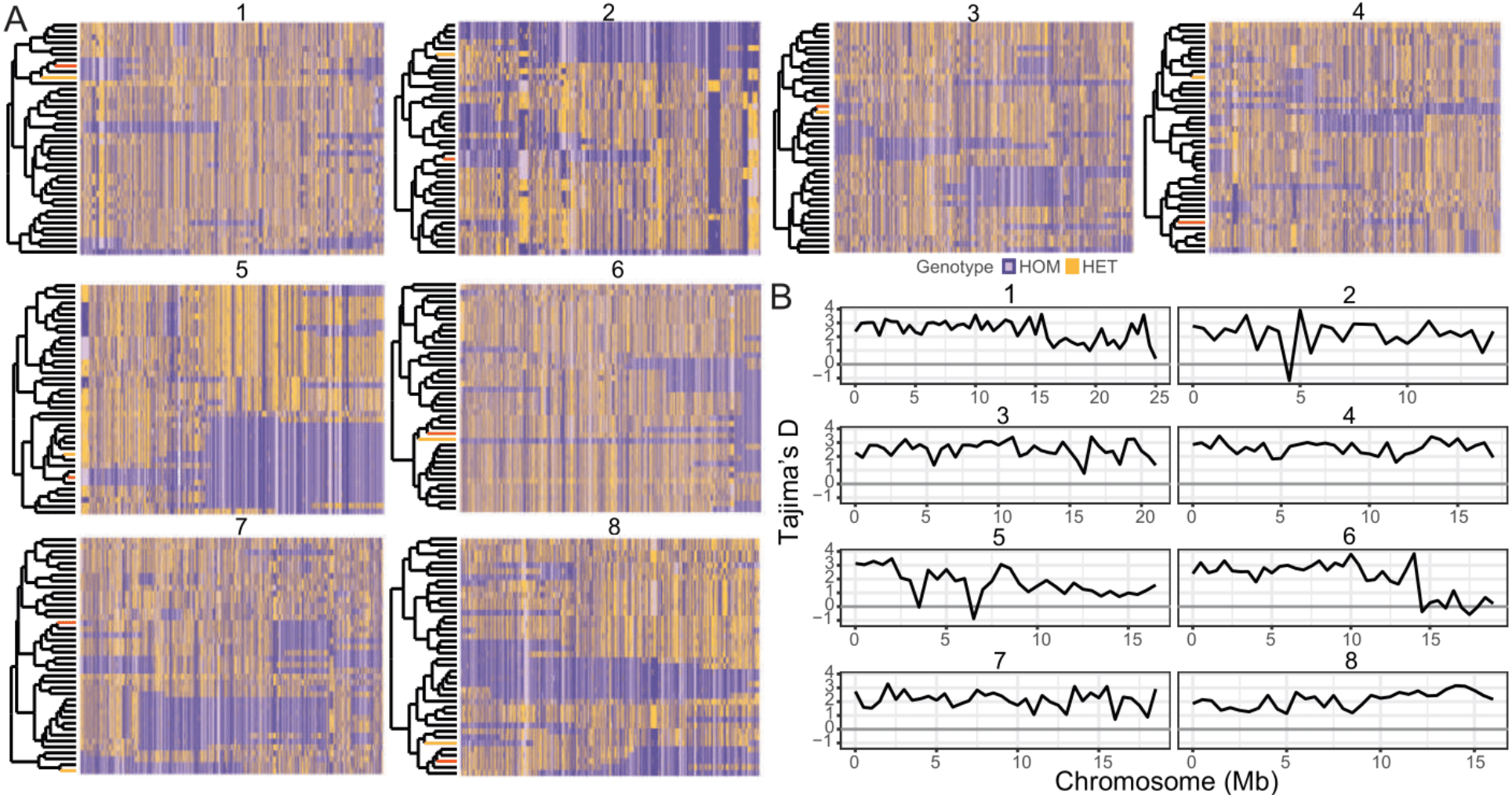
Homozygosity in wild individuals from the Strečno population. **A.** Genome-wide genotype calls from 40 wild plants sampled from the Strečno population. Each chromosome is plotted separately. Individuals clustered by chromosome-wide similarity, with B771 in yellow and B772 in orange. Long homozygous stretches are common among wild individuals. **B.** Genome-wide Tajima’s D values. Positive values are found across most of the genome, hinting at the presence of multiple alleles found at intermediate frequencies in the population. Related to Fig 6.

